# Engineering of a graphene oxide-based two-dimensional platform for immune activation and modulation

**DOI:** 10.1101/2023.08.22.553542

**Authors:** Despoina Despotopoulou, Maria Stylianou, Luis Miguel Arellano, Thomas Kisby, Neus Lozano, Kostas Kostarelos

## Abstract

Nanoscale-based tools for immunomodulation are expected to offer a rich battery of options for more targeted and safer approaches to achieve clinically effective manipulation of the local and systemic immune environment. In this study, we aimed to design nanoscale constructs based on graphene oxide (GO) nanosheets as platform carriers for the TLR7/8 agonist Resiquimod (R848). The non-covalent complexation of R848 molecules on the GO surface resulted in stable complexes by preserving their biological activity. The physicochemical properties, molecular quantification, as well as the overall performance of the complex were systematically investigated. We hypothesized the formation of GO:drug nano-constructs with strong colloidal stability over time, due to the strong π-π interactions between the R848 molecules and the GO surface, and identified that R848 loading efficiency consistently ranged around 75% (of starting molecules), quantified by HPLC and UV-Vis. The 2D morphology of the thin nanosheets was retained after complexation, determined by various (AFM and SEM) microscopic techniques. Based on the surface physicochemical characterization of the complexes by Raman, FTIR, XPS, and XRD, the formation of non-covalent interactions among the GO surface and the R848 molecules was confirmed. Most importantly, GO:R848 complexes did not compromise the biological activity of R848, and effectively activated macrophages *in vitro*. Collectively, this study demonstrates that thin GO sheets can act as platforms for the non-covalent association with small TLR7/8 agonist molecules, forming stable and highly reproducible complexes, that could be exploited as effective immunomodulatory agents.

## 1. Introduction

Immunomodulation is the upregulation or downregulation of the immune responses towards a balanced readjustment of the system after being exposed to autoimmune disorders, infectious diseases, or cancer.[1–4] However, this immune alteration for re-establishing homeostasis is proven to be effective only for a fraction of patients.[5,6] Among the main limitations of this approach are the complexity and heterogeneity of the immunomodulation mechanisms as well as the unexplored dynamic molecular interactions occurring in the immune network.[7–9]

An encouraging advance to overcome this reality introduces novel nanoscale-based immunomodulation platforms attempting to offer specificity and durability of the therapy.[10,11] Among the main advantages of these platforms are the improved pharmacokinetics and tissue distribution of immunomodulating agents and enhanced cellular uptake. A comprehensive understanding of nanomaterials’ physicochemical features is necessary for the design of higher precision therapies aiming at specific tissue or cell type targeting or presentation. Additionally, the sustainability of the effects can be improved as the immunotherapeutic agents may be protected from degradation and slowly released from the nanocarriers.[12–15]

Resiquimod (R848), a second-generation and more potent derivative of FDA-approved imiquimod, belongs to the imidazoquinoline family.[16] Imidazoquinolines are low molecular weight organic compounds with significant antitumour and antiviral features. These features result from their naturally derived, strong immunostimulant properties as they are part of the TLR agonists group.[17,18] R848 is a double TLR agonist (TLR-7, TLR-8) and binds to specific receptors that are located in the endosome. The mechanism of action involves the activation of pro-inflammatory signaling cascades that result in the secretion of cytokines.[19,20]

Imiquimod is clinically relevant to the treatment of several skin malignancies or genital warts. On the other hand, although, several clinical trials explored the use of R848 as a vaccine adjuvant or as a single antitumour agent against melanoma, cutaneous T-Cell lymphoma or actinic keratosis, its FDA-approval has not yet arrived.[20–22] The clinical hindrance for R848 approval is linked to the short blood circulation half-life and more importantly its high off-target toxicity.[23,24] Therefore its performance and safety profile should be improved by using alternative and more targeted delivery strategies.[25,26] Nanomaterials hold great promise in achieving this through encapsulation or surface binding of the specific agonist, to reduce its systemic toxicity while maintaining its biological activity and release patterns.[27,28]

In this frame, 2D materials provide a large surface area that could enable the facile and robust surface adsorption of immunomodulators.[29] Graphene oxide (GO) nanosheets offer additionally a variety of functionalities providing different type of interactions with the desired molecules, good colloidal stability and high dispersibility in biological fluids.[30,31] Well-described and controlled cytoxicity profiles on interaction with a variety of cell types along with the reported biocompatibility and biodegradability are promising for the potential adoption of GO as a carrier for immunotherapeutic agents.[31–33]

In this study, we aimed to take advantage of the above unique combination of properties that GO nanosheets could offer as a carrier and presentation platform of R848 through a facile and highly reproducible non-covalent complexation. We hypothesized that GO and R848 will interact strongly and robustly through π-π interactions and other surface interactions that will lead to stable nano-constructs with immuno-activating capacity. Here, we report the protocol for the preparation of such GO:R848 nanohybrids, along with their thorough structural and surface characterization. Finally, the biological activity of the GO:R848 complexes is demonstrated using a primary macrophage model to demonstrate a proof-of-concept immunoactivation capacity.

## 2. Results

### 2.1. GO nanosheet complex formation with R848

In this study, the starting materials used for the preparation of the GO:R848 complexes were the endotoxin free GO, produced by our group, and the, commercially available, synthetic molecule R848 (**Figure 1A**). The GO material was prepared following the modified Hummer’s method under endotoxin free conditions. The resulting GO nanosheets contained less than 2% of impurities. Also, their average size was below 450 nm and their thickness 1 nm, as found by Atomic Force Microscopy (AFM) and Scanning Electron Microscopy (SEM) (**Table S1**). The lateral dimensions and thickness of the GO materials were selected for optimal interaction with the immune cellular component and the maximum surface area available for R848 loading.[34] Regarding the immunomodulatory agent, R848 is a synthetic tricyclic organic molecule with a molecular weight of 350.8 g/mol. Its chemical structure consists of an imidazoquinolinamine with 2 important substitute groups: an ethoxy methyl and a methyl propanol group. For our studies, the hydrochloride analogue of R848 was chosen, as only this form could be soluble in aqueous solution (1 mg/mL in water).

**Figure 1.**
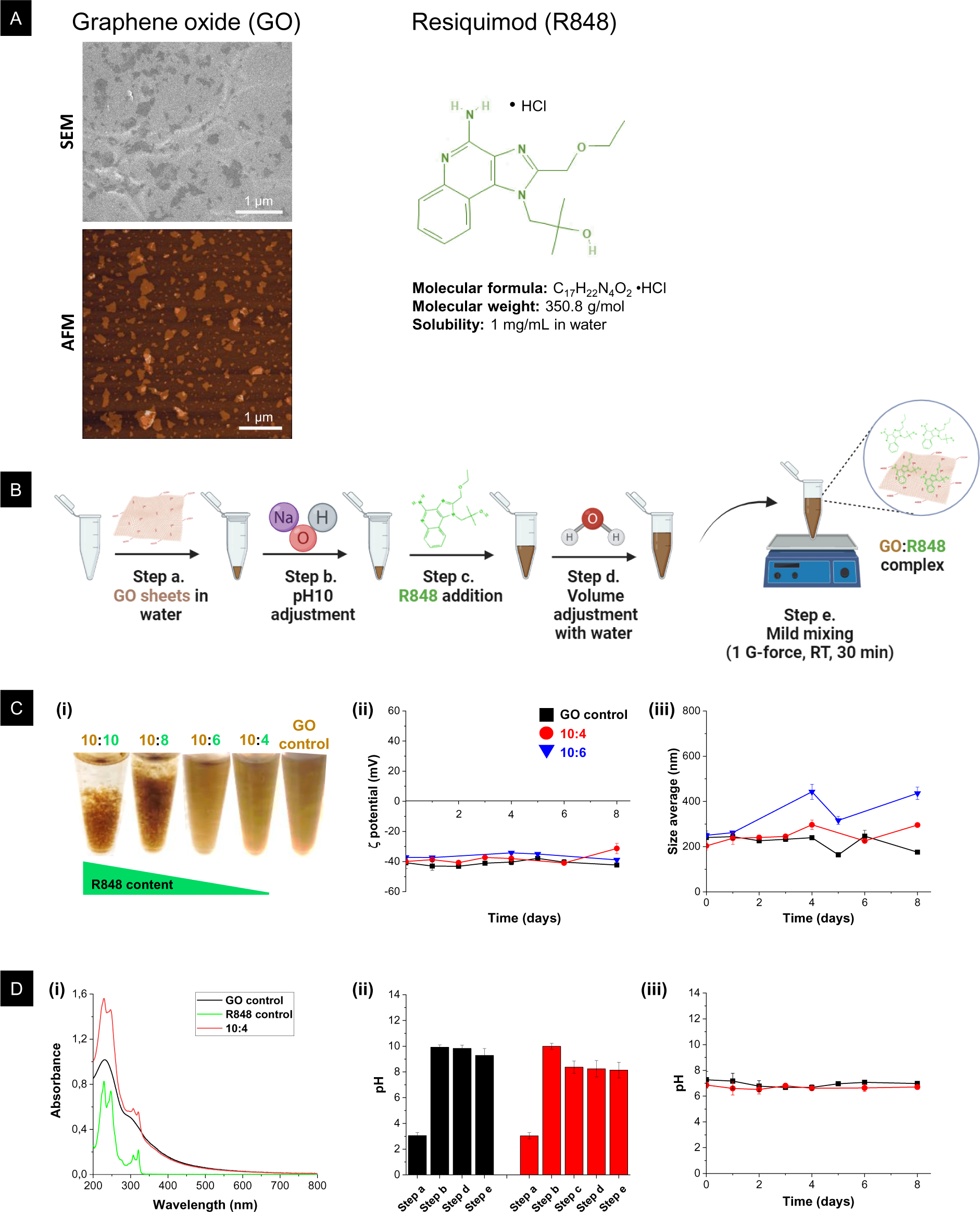
Non-covalent complexation of GO:R848 and construct stability. **(A)** Component material analysis of the GO:R848 system. GO nanosheets synthesized with the modified Hummeŕs method shown by AFM and SEM (scale bars 1μm) and chemical structure and molecular characteristics of R848 molecules. **(B)** Schematic of the non-covalent complexation between GO nanosheets and R848 molecules following four sequential steps of reagent addition and a final incubation/mixing step. Note that at the end of this protocol both adsorbed and unbound R848 molecules will be present. **(C)** Selection of optimal mass (weight) ratio for GO:R848 complex. **(i)** Visual aspect of GO:R848 complexes at different mass (weight) ratios ranging from 10:10 to 10:4, in comparison to GO control. GO control underwent the same protocol but without the addition of R848; **(ii)** Average particle surface charge (ζ-potential) of two GO:R848 complexes at mass ratios 10:6 (blue) and 10:4 (red) in comparison to GO control (black); **(iii)** Mean particle size data by DLS over 8 days at room temperature. Triplicate measurements of at least n=2 sample replicates are shown. (GO (20 µg/mL); GO: R848 (20 µg/mL: 8 µg/mL and 20 µg/mL: 12 µg/mL)). **(D)** Monitoring of GO:R848 complex formation. **(i)** UV-Vis spectroscopic signal of R848 (green) and GO control (black) compared to the optimal GO:R848 (10:4) complex spectrum (red) at day 0. (GO (20 µg/mL); GO: R848 (20 µg/mL: 8 µg/mL); and R848 (8 µg/mL)); **(ii)** pH monitoring for GO control (black) and for the GO:R848 complex 10:4 (red) across the different steps of complexation (a-e), as shown in (B); **(iii)** pH variation over 8 days at room temperature for GO control and GO:R848 complex (10:4); Data expressed as mean±SD of at least n=2 sample replicates. Schematics created with BioRender.com

Complexes between GO nanosheets and R848 molecules were formed in an aqueous solution by moderate stirring, as depicted in **Figure 1B**. Different mass ratios of GO to R848 were tested: 10:10, 10:8, 10:6, and 10:4. The GO:R848 complex at 10:10 and 10:8 (wt:wt) were colloidally unstable upon mixing of the two components, forming a GO aggregate (**Figure 1C(i)**). On the contrary, complexes with ratios 10:6 and 10:4 (wt:wt) visually formed a stable suspension. To establish the optimal GO:R848 mass ratio, complexes at 10:6 and 10:4 were studied by DLS (**Figure 1C(ii) and Figure 1C(iii)**). No significant changes regarding the ζ-potential were observed for these ratios. Nevertheless, a considerable increase in size distribution was detected at the 10:6 ratio (at day 4). These results confirmed a higher colloidal instability in comparison to the 10:4 complex, which was selected as the optimal ratio.

The selected GO:R848 complex (10:4) was carefully monitored since the moment of preparation and over 8 days. At day 0, the complex formation was confirmed spectroscopically by UV-Vis (**Figure 1D(i) and Figure S1**). GO control showed the characteristic peak at 230 nm corresponding to the π-π* transitions of C=C, as well as a shoulder at 300 nm due to n-π* electronic transitions of C=O.[35] On the other hand, the observed adsorption maxima for R848 control was at 220 and 254 nm with other minor peaks at 310 and 320 nm. The fingerprint features of both pure GO and R848 controls were present in the spectrum of the complex, with a notable increase in the absorbance intensity of GO due to the presence of R848. Moreover, pH monitoring of the complex was followed in each preparation step (**Figure 1D(ii))** and over 8 days (**Figure 1D(iii))**. This monitoring was important as the final pH should fall into the physiological range (7-7.4) for biological applications as well as for ensuring electrostatic stabilization of GO sheets.[36] Low pH can cause aggregation driven by the protonation of the carboxyl groups.[37] For this reason, the pH of GO sheets was modified from pH 3 to pH 10. This adjustment in a basic pH was necessary, since the subsequent addition of acidic R848 molecules would have reduced the pH out of the physiological range otherwise.

### 2.2. Purification of GO:R848 complex and quantification of bound R848 molecules

Following optimisation of the suitable ratio to form the GO:R848 complex, a 4-step purification process was developed, using ultracentrifugation with 100 kDa membranes for the removal of the unbound R848 molecules (**Figure 2A**).

**Figure 2.**
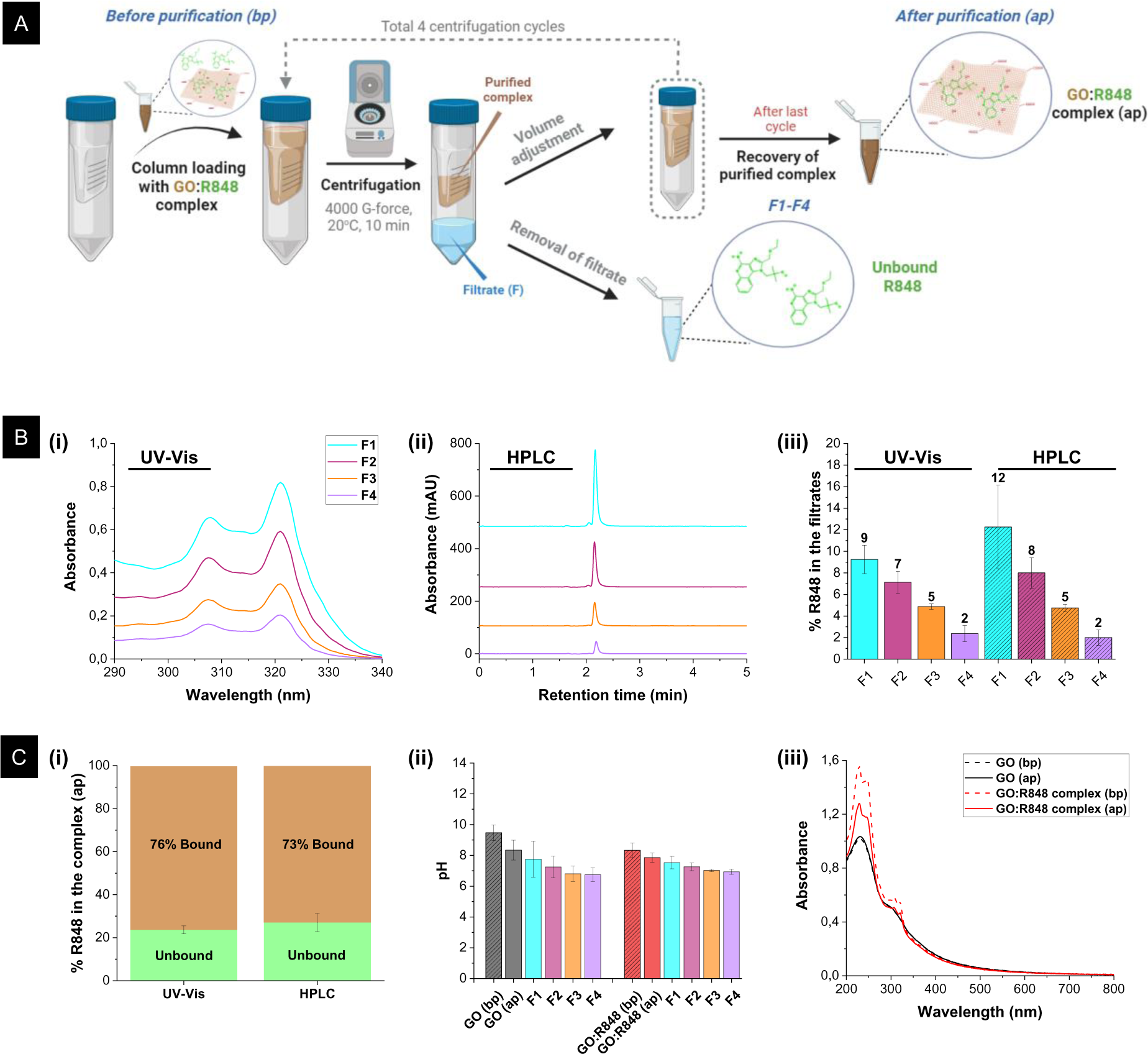
Purification and quantification of bound R848 in the GO:R848 complex. **(A)** Schematic depiction of the purification protocol by using column ultracentrifugation. The final GO:R848 complex after purification (‘GO:R848 (ap)’) was further used for the full physicochemical characterization, while the filtrates F1-F4 (containing any unbound R848 molecules) were used for quantification by UV-Vis and HPLC spectroscopy. **(B)** Quantification of unbound R848 in the GO:R848 (10:4) complex. **(i)** UV-Vis spectrum of GO:R848 filtrates F1-F4 (unbound R848) in the range of 290-340 nm; **(ii)** HPLC chromatogram of GO:R848 filtrates F1-F4; **(iii)** % percentage of R848 after GO:R848 complex purification in the filtrates F1-F4 assessed by UV-Vis and HPLC methods. The results (generated by applying the formula (1) shown in the experimental section) are expressed as mean±SD of n=3 replicates. **(C)** Effect of purification method on the GO:R848 (10:4) complex. **(i)** Percent (% of originally added) of unbound R848 (green) and bound R848 to the GO surface (brown) as measured by both quantification techniques (UV-Vis and HPLC). The results (generated by applying the formula (1)) are expressed as mean±SD of n=3 replicates; **(ii)** pH monitoring at different steps of purification: GO control (black) and GO:R848 complex (red) before (bp) and after (ap) purification appeared in dashed and solid bars respectively, and their corresponding filtrates (F1-F4) during the purification steps. Data are expressed as mean±SD of at least n=3; **(iii)** UV-Vis spectra of the GO control (black) and the GO:R848 complex (red) capturing the signal before (bp) (dashed line) and after (ap) purification (solid line) in the range 200-800 nm (GO (20 µg/mL)); GO: R848 (20 µg/mL: 8 µg/mL)). Schematics created with BioRender.com.

Before purification of the GO:R848 complexes, the GO and R848 controls were tested for the validity of purification procedure, therefore for assessing if a clear separation between GO and the R848 molecules could be achieved. For the GO quantification, the UV-Vis calibration curve at 230 nm was used.[38] For R848, the wavelengths of 320 nm and 254 nm were selected for the UV-Vis and HPLC calibration curve respectively, as depicted in **Figure S1**.[39–41] The comparison of the UV-Vis peak absorbance (at 230 nm) for GO before and after purification indicated the total recovery of GO concentration, suggesting that there was no significant retention or loss of GO in the ultrafiltration membrane. In addition, the UV-Vis absorbance and HPLC analysis of the GO filtrates (F1-F4) further proved that GO is not passing through the membrane (**Figure S2A**). Regarding the R848 control, it was found by UV-Vis spectroscopy, that around 99% of the initial R848 crossed the membrane in the initial two purification cycles and into the filtrates (95% in F1 and 4% in F2). Moreover, HPLC results validated these findings (94% in F1 and 3% in F2) with a final recovery of 97% (**Figure S2B**). Overall, 98-100% GO could be quantified by UV-Vis in the purified fraction (material on the filter), and 97-99% of R848 could be quantified by both UV-Vis and HPLC in the filtrates, as shown in **Figure S2C**, indicating the suitability of the method for quantification of the amount drug molecules complexed onto the GO.

Subsequently, the loading capacity of R848 onto the GO nanosheets was studied by UV-Vis spectroscopy and HPLC, by measuring the unbound R848 molecules washed through the membrane and collected into the filtrates. The UV-Vis absorption peak intensity for R848 in the filtrates during the purification cycles of the GO:R848 complexes, indicated a descending trend of 9%, 7%, 5% and 2% of R848 in the filtrates F1-F4 respectively. By HPLC these were found to be 12%, 8%, 5% and 2%. Both techniques illustrated the necessity for four centrifugation cycles to achieve removal of unbound R848 (**Figure 2B**). The bound R848, calculated from the measured unbound R848, was found to be between around 73-76% by HPLC and UV-Vis (**Figure 2C(i)**). Overall, it was found that for each 1000µg of GO in the complex, the actual amount of R848 complexing was ca. 300µg (855 µM).

In addition, the effect of the purification process onto the GO:R848 complex was explored. Following the ultrafiltration process, the pH of GO and GO:R848 is reduced and remained neutral as was intended due to the removal of any excess sodium hydroxide in the filtrates (**Figure 2C(ii)**). Also, the purified GO:R848 complex still showed the characteristic peaks of both individual components with an additional reduction in the absorbance intensity in comparison with the unpurified complex. The purified GO:R848 complex absorbance intensity was overall reduced due to the removal of the unbound R848 moieties (**Figure 2C(iii)**).

### 2.3. Structural characterization of the GO:R848 complexes

The structural features and morphology of the GO:R848 complexes were studied by AFM and SEM. Both were compared to the unmodified GO control. The AFM images showed that the flat morphology and thickness profile of the GO nanosheets was retained after R848 complexation, indicated that the drug loading was not provoking any stacking effect (**Figure 3A**). Interestingly, the nanosheet cross section analysis revealed an uniform height of 0.9 to 1.1 nm, suggesting a complex that consists of 1 layer.[42] Moreover, the SEM images showed a smooth lamellae surface, an arrangement which is consistent with the preservation of the typical polygonal morphology of GO nanosheets with a thin layer structure after complexation with R848 (**Figure 3B**). Additionally, lateral size distribution analysis of AFM and SEM data demonstrated that the size range of the complexed nanosheets were very similar to those of GO alone (**Figure S3**). It is worth mentioning that the size reported by SEM and AFM techniques differs from the value that was measured by DLS during the complexes preparation, due to the limitation of the DLS technique in the precise measurement of 2D planar sheets.[43]

**Figure 3.**
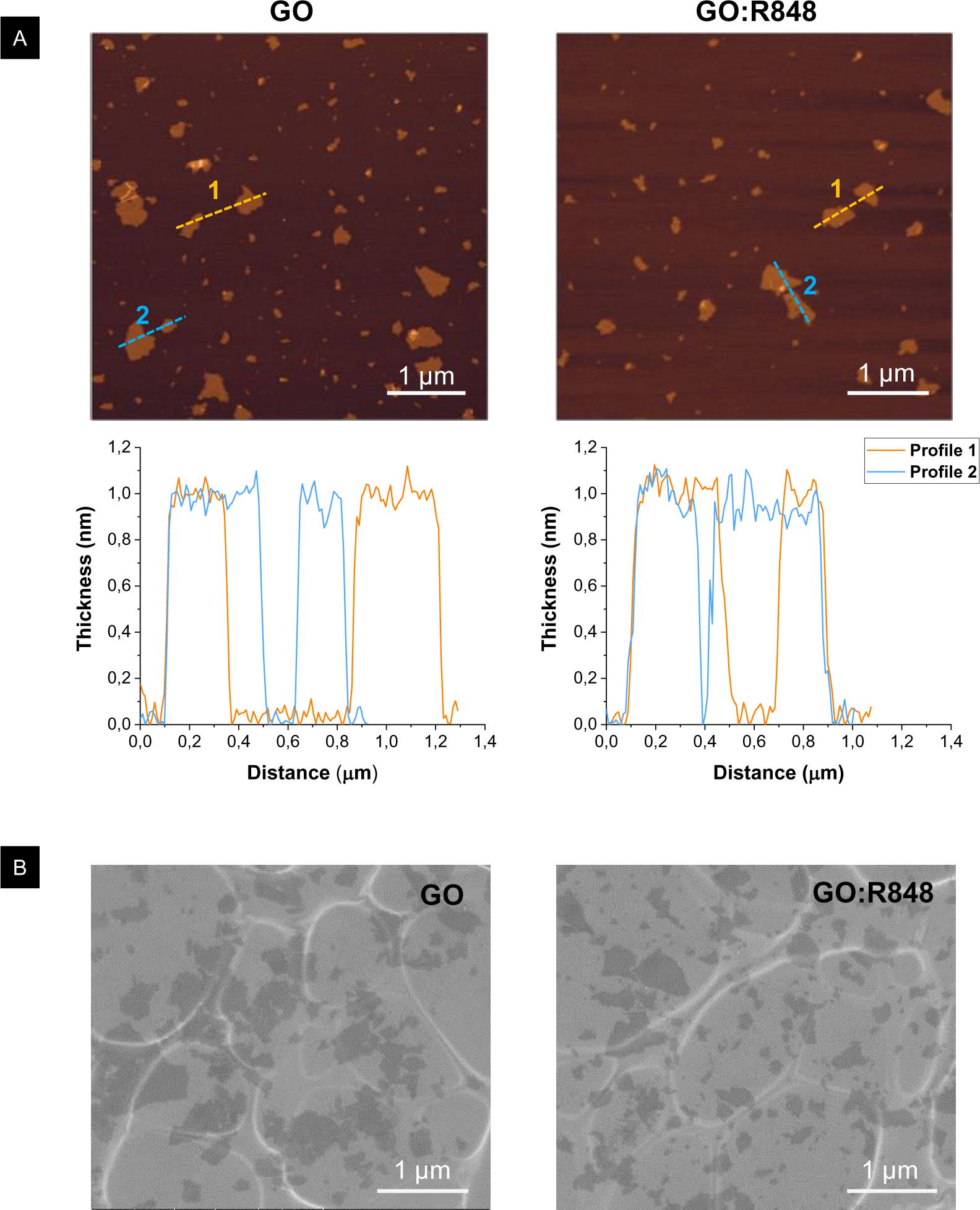
Structural characterization of GO:R848 complex after purification (ap) by AFM and SEM. **(A)** AFM height images with corresponding nanosheet cross-section and thickness graphs (shown underneath) of: **(i)** GO control; and **(ii)** GO:R848 complex. **(B)** SEM micrographs of: **(i)** GO control; and **(ii)** GO:R848 complex. In all images scale bars are 1μm.

### 2.4. Atomic and elemental surface characterization of GO:R848 complexes

Detailed atomic-level surface characterization was performed using a battery of spectroscopic techniques, namely Raman spectroscopy, Fourier transform infrared (FTIR), X-ray photoemission spectroscopy (XPS) and X-Ray diffraction (XRD).

The Raman data were collected using a 633nm laser, and the structural disorder ratio (I_D_/I_G_) was calculated (**Figure 4A**). It is well known that GO material alone displays typically two peaks: a D band assigned to disordered sp^3^-based defects, and a G band ascribed to sp^2^-hybridized carbon at 1327 cm^-1^ and 1601 cm^-1^ respectively.[44,45] When complexation with the R848 molecules was performed, it was evident that the spectra for the GO:R848 complex contained contributions mainly from the GO component. However, new spectroscopic features at around 995, 1473, and 1528 cm^-1^ ascribed to the imidazoquinoline unit were clearly detected.[46] Turning our attention to the G band, this mode (1589 cm^-1^) was downshifted by 12 cm^-1^ with respect to GO due to the n-doping effect of the R848 as observed in the literature with different electroactive units.[47] Finally, the comparison between the I_D_/I_G_ ratios can provide further information on the materials structure. The I_D_/I_G_ ratio is related to the GO surface oxidation degree and allows for estimation of the degree of defects on the graphitic lattice.[48] In GO:R848 the I_D_/I_G_ ratio was found 1.11, slightly lower than for GO starting material (1.24), which may be due to the surface interactions with the R848 and the slight distortion of the GO bands. However, further techniques were explored to affirm these interactions and their effect on the GO lattice.

**Figure 4.**
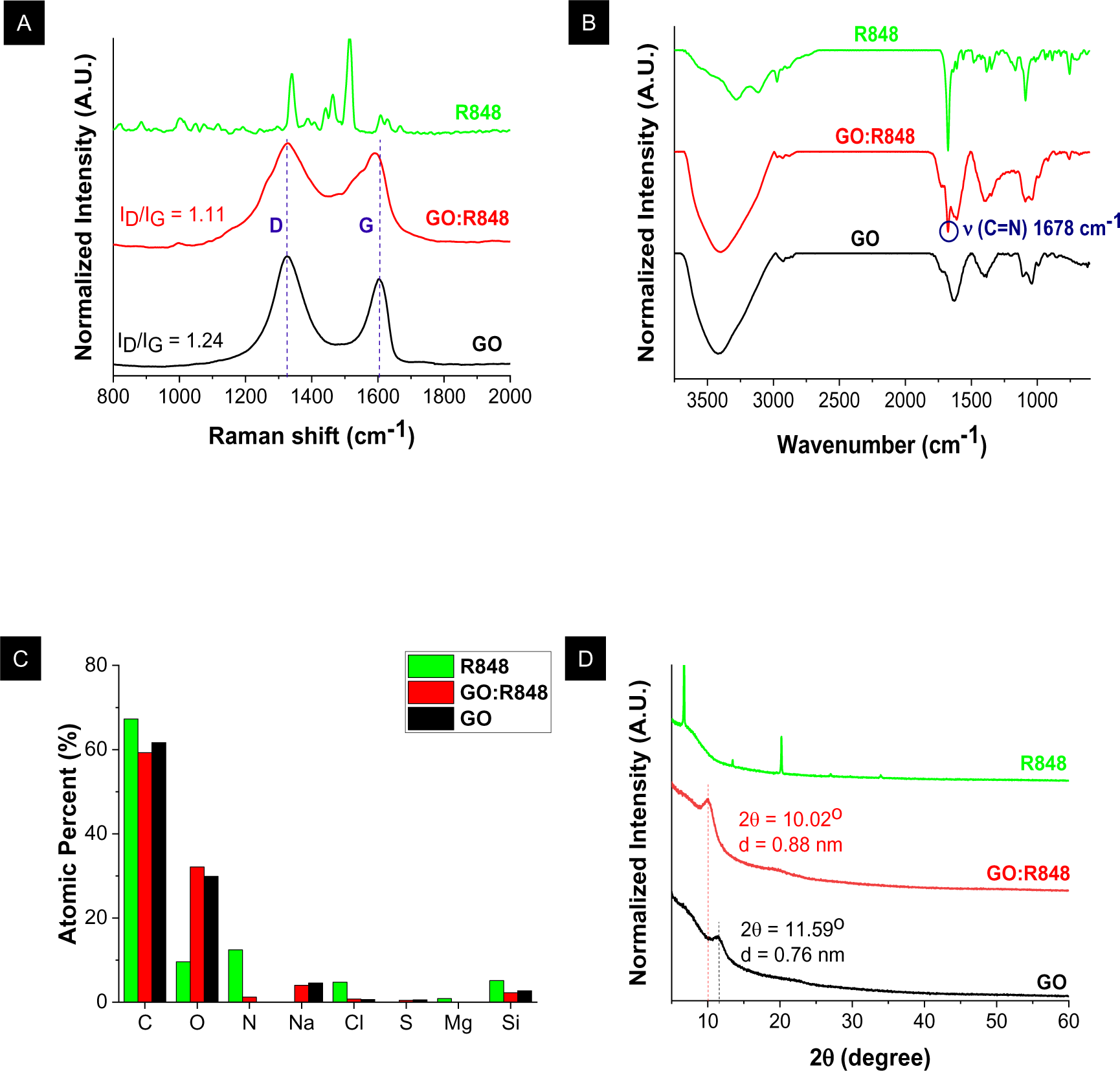
Atomic-level and elemental characterization of GO:R848 complex after purification (ap) by surface spectroscopy. For comparative reasons, the data for GO alone (black) and R848 alone (green) controls are also presented. **(A)** Raman spectrum with the I_D_ and I_G_ bands underlined. The I_D_/I_G_ ratios are noted. **(B)** FTIR spectrum with the contribution of the characteristic surface functionality included. **(C)** Total contribution of various functional groups in relative atomic percentages (%) as measured by XPS. Note that Si contribution derives from the Si substrate used during measurement. **(D)** XRD spectrum with the corresponding interlayer distance highlighted. Peak intensities were generally normalized by dividing with the highest value.

Successful complexation of R848 onto GO was also confirmed by interpreting the solid state FTIR spectra of the GO:R848 complex compared to the GO control (**Figure 4B**). The infrared spectrum of GO showed the characteristic vibration modes corresponding to the following functional groups; v(C-H) at 2850 cm^-1^, v(C=O) at 1726cm^-1^, v(C=C) at 1625cm^-1^, v(O-H) at 1400cm^-1^ and v(C-O) at 1046cm^-1^.[38,49] The GO:R848 complex FTIR data consisted of contributions from both the GO and R848 components. The prominent peak at 1678cm^-1^ due to stretching vibration C=N of the imidazole ring, as well as a peak of weak intensity at 750cm^-1^ due to the chlorine content (since the R848 is a chlorine salt), confirmed association of R848 onto the GO surface.[50–52] The absence of new vibrational bands in the GO:R848 apart from the individual GO and R848 contributions, affirmed the physical association of the molecule without the formation of any chemical bonds. Finally, the absence of many vibrational bands of R848 in the GO:R848 spectrum may indicate a strong interaction with the GO surface restricting the vibration modes of the organic moieties, as has been previously reported for other molecules with aromatic rings.[53,54]

Additional evidence for GO surface modification was obtained from X-ray photoelectron spectroscopy (XPS) by measuring changes in the elemental composition at the surface of those nanosheets. In general, survey spectra for GO, R848 and GO:R848 exhibited signals corresponding to electrons from C_1s_, O_1s_ and N_1s_ core-level (**Figure S4A(i)**). The C_1s_ and O_1s_ peaks appeared in the binding energies of 285eV and 531eV respectively. The presence of the above mentioned peaks in the same eV both for the starting material and the GO:R848 confirmed that complexation of R848 onto the GO surface preserved the chemical composition and properties of the starting material. After loading R848, the existence of a novel N_1s_ contribution at a binding energy of around 400 eV ascribed to the nitrogen content of R848 (not present at all in GO), indicated the occurrence of complexation (**Figure S4A(ii) and Figure S4B**). Quantitative data obtained from XPS measurements are presented in **Figure 4C**. In all samples, the detected Si is attributed to the underlying supporting Si substrate used for the measurements. Also, the detected Cl in R848 was around 5%, due to the hydrochloride analogue that was used. Trace elements around 1% of total atomic composition (Cl, S, Mg) may come from impurities during the synthetic procedure. Both for GO and GO:R848, around 60% was C and 30% O, the main two elements of the GO sheets, and around 4% Na, due to the neutralization of the GO sheets with sodium hydroxide. Interestingly, a small amount of N (1%) indicated the presence of R848 in the GO sheets.

The XRD method (**Figure 4D**) was used to further interrogate the formation of GO:R848 complexes by comparing the interlayer distance between the nanosheets, as well as possible structural disruptions in the case of the GO:R848 complex. In the XRD diffraction pattern of R848, intense crystalline peaks appeared around 7° and 20° as previously described for imidazoquinoline derivatives.[55] For the GO control, the characteristic peak was obtained as a broad and diffuse signal at 2θ=11.59⁰ with basal spacing of 0.76 nm, as calculated by Braggs Law.[56] This suggests that GO consists of an amorphous structure which is also retained after R848 complexation. For the GO:R848 complex, the GO peak is shifted to lower 2θ=10.02⁰, which corresponds to a slightly higher basal spacing of 0.88 nm and can be attributed to intercalation of the therapeutic modality and a possible re-arrangement of the GO nanosheets.[57]

### 2.5. Colloidal stability of GO:R848 complexes

The colloidal stability of nanomaterial suspensions is an important factor that can be directly influenced by the interactions between the GO nanosheet surface and the nature of the molecule used to form the complex. To evaluate the long-term stability for both the GO and GO:R848 complex aqueous suspensions, their pH, mean particle diameter and ζ-potential were monitored over 2 months. The pH of the GO control and the GO:R848 complexes remained stable (pH ̴ 7-8), the size distribution curves (by DLS) ranged between 200-300 nm with a good Gaussian distribution and the ζ-potential fluctuated between −35 to −45 mV, all suggesting maintenance of colloidal stability (**Figure S5A and Figure S5B**). Interestingly, lack of any R848 detachment from the GO surface within a period of 2 months under storage in dark conditions and at room temperature indicated strong π-π interactions between GO and R848 (**Figure S5C**). The long-term morphology of the nanosheets was analysed by AFM and SEM (**Figure S5D and Figure S5E**). No nanosheet stacking or severe morphological changes were observed both for the GO control and the GO:R848 complexes. The thickness of the sheets in both suspensions was found stable between 1-1.2 nm, indicating the high stability of the materials after 2 months. The size distribution data (**Figure S3**) suggested that the nanosheets were able to preserve their initial structural characteristics and were comparable between the GO alone and its complex with R848.

Lastly, prior to the biological studies, it was necessary to assess the colloidal properties of GO and GO:R848 in the cell culture media (DMEM + 10% FBS) for 24 hours. Visually, some clusters formed at 24 h possibly due to the nanosheetś protein coating (**Figure S6A**). Both at 0h and 8 h, as shown by AFM, the protein association on the GO surface can be observed. However, the typical polygonal morphology of the flakes was retained in all cases, suggesting colloidally stable nanosuspensions that could be further used for the *in vitro* studies **(Figure S6B and Figure S6C).**

### 2.6. Biological activity of GO:R848 using primary macrophages

After the formation of the GO:R848 complex, the structural characteristics and the colloidal stability of the suspensions were proved and described in the previous sections, one of the final aspects for investigation was the complex bioactivity. The main question needed to be answered was whether the surface association of R848 onto the GO surface, was compromising the biological potency of the immunomodulator. In order to elucidate this, the activity of GO:R848 was investigated in an *in vitro* setting. Murine bone marrow derived macrophages (BMDMs) were exposed to the complex for 2 hours and this interaction was evaluated. GO is known to possess distinct autofluorescence properties enabling tracking and imaging of the interaction with live cells.[58] Following exposure of BMDMs, both GO and GO:R848 treated cells showed and increased auto fluorescence which co-registered with the presence of black particles (attributed to GO) specifically in these cells confirming the interaction of both GO and GO:R848 with the cells (**Figure 5A and Figure S7**). Following, the biological activity of the complex was investigated by analysing the immunoactivation of BMDMs by expression of an activation marker CD80 via flow cytometry. It was demonstrated that GO:R848 could increase the percentage of macrophages (F480+/CD11b+) expressing CD80 to the same levels as free R848, confirming that the biological activity is maintained (**Figure 5B and Figure S8**). To further validate these results, the production of TNF-α, a key pro-inflammatory cytokine that is released upon activation of macrophages was also assesed. TNF-α levels were significantly upregulated in GO:R848 treated cells compared to the controls including free R848 (**Figure 5C**). Overall these results confirmed that GO:R848 can act as a suitable platform for the delivery of R848 to immune cells while maintaining the immunostimulatory activity, therefore highlighting the potential for future use as an immunotherapeutic nano-construct.

**Figure 5.**
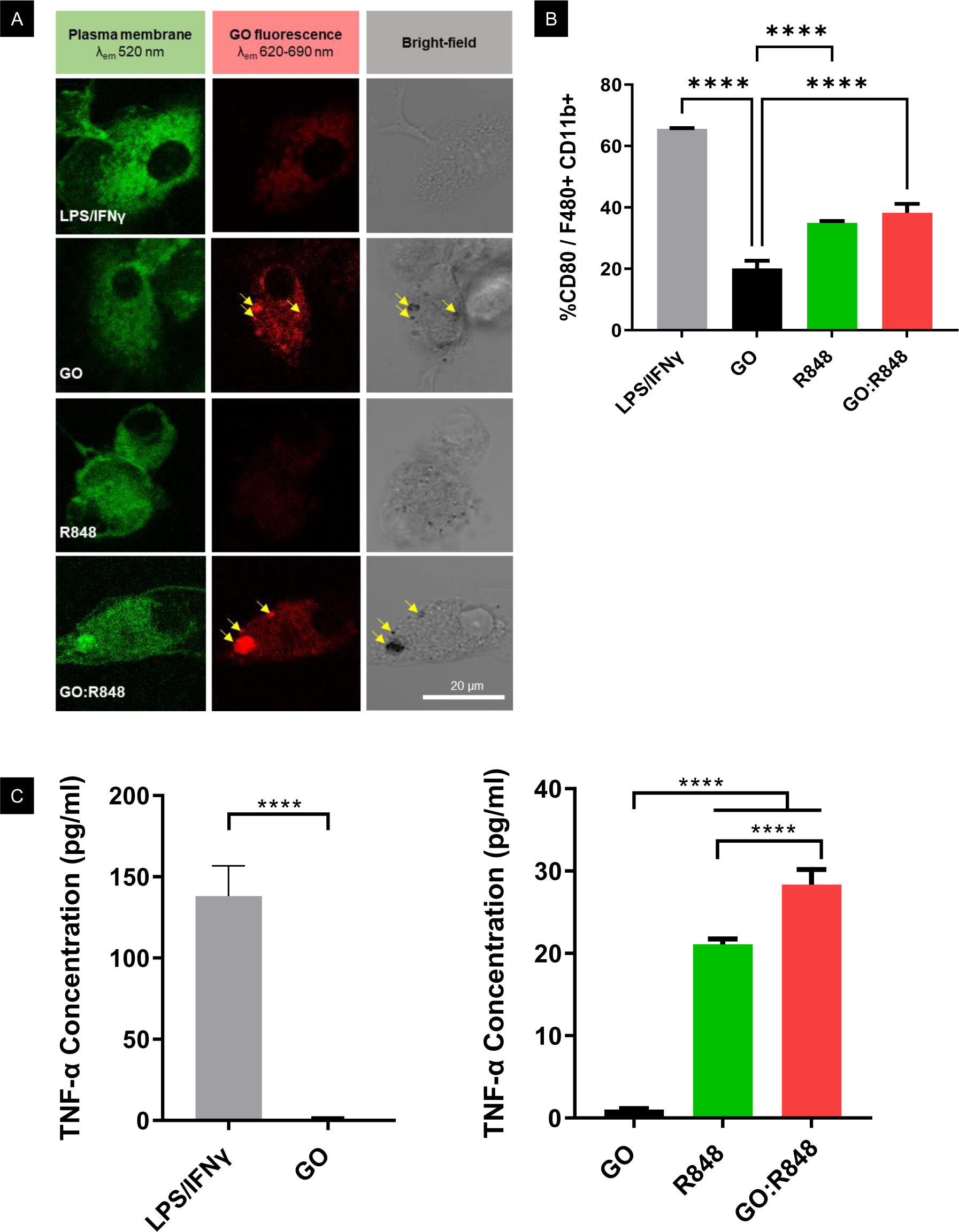
Biological activity of the GO:R848 complex demonstrated by effective activation of primary macrophages *in vitro*. **(A)** Interaction of GO and GO:R848 complex with primary bone marrow derived macrophage (BMDMs) membranes, 2hr post treatment. Cells were stained with green plasma membrane dye. Bright field confirmed the presence of GO (black particles in bright field). GO autofluorescence (red channel) was used to detect GO-BMDMs co-localisation and distinguish it from cell auto-fluorescence. All images were obtained with confocal microscopy at 40x magnification, zoom factor 2, scale bar 20μm. **(B)** Percentage of CD80 (activation marker) out of F480+ CD11b+ cells, 24hr post-treatment with LPS (100 ng/mL); IFNγ (20 ng/mL), GO (10 µg/mL), GO:R848 (10 µg/mL: 4 µg/mL) and R848 (4 µg/mL); n= 3 biological replicates per condition. **(C)** TNF-a expression (pg/ml) of BMDMs supernatant, 24 hr post-treatment as mentioned above; n= 6 biological replicates per condition. Data is presented as mean±S.D. Ordinary one-way ANOVA, Tukey’s multiple comparisons test (*p ≤ 0.05, ** p ≤ 0.01, ***p ≤ 0.001, ****p ≤ 0.0001).

## 3. Discussion

In this study, a robust protocol for the non-covalent complexation and purification of GO with the immunomodulating molecule R848 was presented. The GO:R848 complexes showed a drug loading capacity of 75% and a negative surface charge that prevented aggregation.[59] It was shown that the complexed R848 did not bear any significant structural impact on the thickness and lateral dimension of the ensuing nanosheets. Detailed characterisation of the complexes using an array of different techniques strongly confirmed the presence and strong interaction of R848 molecules onto the GO surface. Long-term stability studies demonstrated that the complex remained a colloidally stable suspension, without any morphological changes or R848 detachment from the GO surface, further indicating the strong physical interactions between the two complex components. Most importantly, the GO:R848 complex was shown to preserve the immunostimulatory activity of R848 in primary macrophages cultures.

Up to now, limited studies have been published regarding the formation of two dimensional GO:R848 complexes. A chemically functionalized GO platform with amino-thiophenol crosslinked to polyethylenimine was used for the combined delivery of plasmid DNA (OVA-encoding) and of the immunomodulator R848.[60] Similarly, Yin et al. designed a hydrogel as cancer nanovaccine that contained GO flakes chemically functionalised with polyethylenimine (PEI) complexing mRNA (OVA-encoding) and the R848 adjuvant.[61] Despite these two reports exploring the use of R848 molecules as vaccine adjuvants, no thorough atomic level characterisation of a GO:R848 complex has been previously presented. Here we attempted to unravel a method to the generation and characterisation of a purified GO:R848 complex using a battery of different structural, surface, elemental and colloidal investigations. It is worth mentioning that throughout these studies, GO alone was used as a control. We intentionally did not introduce any functional groups to the GO surface, hypothesizing that the strong non-covalent interactions between the highly purified and thin GO lattice with the R848 molecules will lead to a stable complex with minimal interference to the surface properties of the nanosheets.

The use of R848 and other immune-modulating molecules in that family has not been greatly explored in combination with 2D materials. Sun et al. used boron nanosheets, prepared by a liquid exfoliation technique from bulk boron, modified with polydopamine and loaded with tumour antigen and R848 in a platform combining immunotherapy and phototherapy.[62] In this report, R848 was also loaded on a polymer-functionalized 2D surface, so clear R848 interactions with GO were not explored. Despite the limited studies using 2D materials, R848 has been successfully encapsulated within a few different types of nanocarriers, including poly(lactic-co-glycolic acid) (PLGA), cyclodextrin, hyaluronic acid polymeric platforms and Au nanoparticles.[41,63–67] Notably, in the majority of the above, non-covalent association of R848 with the nanocarriers was reported to allow for minimal interference with the R848 inherent chemical and biological properties, since minor modifications of its structure can alter significantly its potency.[68] Our approach in forming the GO:R848 complexes non-covalently was along the same lines, along with sufficient complexation efficiency (around 75% of starting R848 molecules) presumably due to the large surface area on the GO lattice available and the strong π-π interactions between the two components. In the case of PLGA or gold nanoparticles, lower entrapment efficiencies have been reported (around 8% and 30% respectively), maybe due to weaker interactions between the R848 molecules and the carrier nanoparticles.[41,67]

On the biological front, concerted efforts need to be made for strategies that overcome the reported systemic toxicity from R848 administration. Among such approaches, the preparation of prodrug formulations, as tocopherol-functionalized R848 loaded into polymeric carriers, or azide-masked R848 have been described.[69,70] Also, smart responsive nanosystems through hydrolysable bonds have been designed for specific release of the R848 in targeted areas (e.g lysosomes).[25,28,71,72] In our approach, emphasis was placed on the adherence and stable complex formation between the nanocarrier and the R848 molecules in order to minimise toxicity risks from free or detached R848. Furthermore, the capability of the GO nanosheets to internalise within immune system components is critical to achieve effective immunomodulatory effects. For example, Li et al. has reported targeting of dendritic cells through mannose-functionalization of the nanoparticles.[73] We had previously demonstrated the inherent immune cell component affinity of the thin GO nanosheets used in this study, able to be internalized by macrophages resident in the tumour microenvironment.[57] Demonstration in this work that such properties can also be preserved for the GO:R848 complex are critical in their exploration as an immunomodulatory platform. Even though most of the previously published work using R848 has been oriented towards cancer immunotherapy via macrophage repolarization, chemoimmunotherapy, or combinations of immunotherapy with phototherapy,[66,74,75] a limited number of reports also describe nanocarrier-based R848 immunomodulation in different applications, such as infections or allergies.[76,77]

Overall, R848 has been previously identified as a potent immunoadjuvant for innate immune cell activation, including macrophages and dendritic cells[78–80]. Delivery and presentation of R848 to these cells types, or to sites where these cell types reside, can improve both immunomodulatory activity and reduce off-target effects. Toward this, we demonstrated that a purified and highly stable GO:R848 nano-construct could interact with and effectively activate macrophages *in vitro*, confirming the preservation of its immunomodulatory capacity. Further investigations are both warranted and necessary to elucidate the mechanisms of GO:R848 activity, as well as to investigate whether these complexes can contribute to improved therapeutic effects in the context of a disease model.

## 4. Conclusion

This study has demonstrated the efficacy of thin GO nanosheets as a carrier platform for the immunomodulator R848 that has been considered very promising in immunotherapy applications. The formation of the GO:R848 complexes was shown by the combination of multiple characterization techniques, indicating the strong interaction between R848 and the GO lattice through non-covalent interactions. The described system is encouraging for further biological investigations due to its chemical and colloidal stability over time and, more importantly, for the preservation of its bioactivity upon complexation. Altogether, this is a simple and highly reproducible method for the complexation of R848 onto GO nanosheets aiming at the generation of flat-shaped nano-constructs with enhanced immunoactivity. Further studies are warranted to determine the *in vivo* immunomodulatory effects of such nano-constructs and their impact on improved therapeutic outcomes.

## 5. Experimental section

### Reagents

Endotoxin-free GO was internally provided by our group and was prepared with a modified Hummers method, as previously described.[38] Resiquimod (R848) was purchased from InvivoGen (sterile, ≥ 95% HPLC purity). Water for injections (WFI) used for the graphene oxide production and complexes preparation was purchased from Gibco. Additional chemicals used were purchased from Sigma-Aldrich (Merck, Spain). Cell culture reagents were purchased from Sigma-Aldrich (Merck, UK) unless stated otherwise. Amicon Ultra-4 100 kDa MWCO Centrifugal Filter units were purchased from Merk-Milipore (UFC810024).

### GO:R848 complex preparation

Firstly, R848 powder was resuspended in water for injection to obtain a stock solution of 1 mg/mL and stored at −20 °C. GO solution was adjusted to a final pH of 10 with 0.1 M sodium hydroxide. The surface association of R848 onto GO was performed by a mild mixing of R848 with the GO sheets in water, at different GO:R848 mass ratios (wt:wt) 10:10, 10:8, 10:6, and 10:4. GO nanosheets at an initial concentration of 1000 µg/mL were mixed with R848 molecules at a certain concentration of 1000, 800, 600, and 400 µg/mL respectively. The overall complexation volume was kept constant at 1 mL by the addition of water. Then, the obtained suspension was incubated under mild shaking conditions (1 G), at room temperature for 30 minutes, followed by 1 h of stabilization. For the controls, free R848 was diluted at a final concentration of 400 µg/mL. GO control, at a concentration of 1000 µg/mL, followed the same protocol described above, without the step of R848 addition.

### GO:R848 complex purification and quantification of bound R848 molecules

For the purification of the complex mixture from any unbound R848 moieties, 100 kDa Amicon Ultra Centrifugal Filter units were used. For that step, the complex 10:4 was centrifuged 4 times at 4000 G, at 20 °C for 10 minutes. After each cycle, the filtrate was collected, and the volume of the purified fraction (on top of the membrane) was adjusted with water for the next round. The same procedure was also followed for the GO control for comparative reasons. Next, the loading amount of R848 on the GO sheets was indirectly calculated by measuring the concentration of unbound R848 present in the collected fractions with UV-Vis spectroscopy and HPLC. The formula used for the calculation of the %bound R848 (drug entrapment efficiency) was the following:

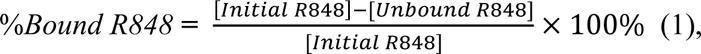

where [Ιnitial R848]: theoretical R848 concentration mixed with GO (in µg/mL) and [Unbound R848]: quantified R848 collected in the filtrates after purification (in µg/mL)

### UV-Vis Spectroscopy

GO and R848 samples were diluted in water at concentrations ranging from 2.5 to 20 µg/mL and from 2 to 40 µg/mL respectively and they were measured using a Hellma QS Quartz cuvette. Absorbance spectra were recorded using the Evolution 201 UV-Vis spectrophotometer (Thermo Scientific) in the range of 200-800 nm at room temperature. The standard curves were obtained for GO at 230nm and for R848 at 320nm in above-mentioned concentrations and showed high linearity with a correlation coefficient (R^2^) of 0.999. Unloaded R848 was calculated by interpolation from the calibration curve. The spectra were analyzed with Origin software (version b9.5.0.193).

### High-Performance Liquid Chromatography

R848 chromatograms were recorded by HPLC system (PerkinElmer Flexar) with a multiwavelength UV-Vis photodetector and a C18 Hypersil BDS column (4.6 x 150 mm, 5 µm, ThermoFisher). The chromatographic analysis was carried out using mobile phase A (0.1 % TFA in water) and mobile phase B (0.1 % TFA in acetonitrile) following an isocratic elution (50:50 v/v) and a flow rate of 1 mL/min. The injection volume was 10 µL. R848 was detected at 254 nm and the retention time was approximately 2.1 ± 0.05 min. The standard curve was linear over the concentration of 5 to 100 µg/mL with a correlation coefficient (R^2^) of 0.999.

### Atomic Force Microscopy

Samples were prepared by covering a cleaved mica surface (Ted Pella) with 20 µL of 0.01 % poly-L-lysine solution (Sigma-Aldrich). After washing with water, 20 µL of GO solution at a concentration of 0.1 mg/mL were drop-casted and washed again with water. For the measurements, the atomic force microscope Asylum MFP-3D (Oxford instrument) was used in air-tapping mode. Silicon probes (Ted Pella) with a resonance frequency of 300 kHz and nominal force 40 N/m were used. AFM images of 5 µm × 5 µm were processed with Gwyddion software (version 2.57).

### Scanning Electron Microscopy

20 µL of samples with GO concentrations of 0.1 mg/mL were dropcasted on grids with Ultrathin C on the Lacey C film (Ted Pella) and the excess of the droplet was removed by blotting. The procedure was repeated 4 times and the samples were dried overnight at room temperature. SEM images were recorded at the ICN2 Electron Microscopy Unit with a Magellan 400L field emission microscope (Oxford instruments) and an Everhart-Thornley detector for secondary electrons. The measurement conditions were 100 pA beam current and 20kV acceleration voltage. The image processing was performed using Image J software (version 1.8.0).

### Raman Spectroscopy

Samples were prepared by drop-casting 20 µL of 0.1 mg/mL GO on top of a glass coverslip and then dried overnight at room temperature. A confocal Raman microscope (Witec) under laser excitation of 633 nm and 600 g/nm gradient was used for the recording of the measurements. Power of 1 mW for 10 s was used to irradiate the sample and for the focusing an objective lens of 50 x magnification was selected. The spectra baseline was corrected and the I_D_/I_G_ ratios were calculated on the maximum of the D and G bands of each sample.

### Fourier Transform Infrared Spectroscopy

Samples were prepared on potassium bromide via a casting process. The spectra were collected at the ICN2 Molecular Spectroscopy and Optical Microscopy Facility with a Tensor 27 FT-IR spectrometer (Bruker). A resolution of 4 cm^-1^, a scan range of 3750 to 600 cm^-1^ and a baseline correction treatment were selected.

### X-ray Photoemission Spectroscopy

20 µL of the sample were drop casted several times until a thin film was formed on top of a 5×5 mm Si wafer (Ted Pella). A Phoibos 150 (SPECS, GmbH) electron spectrometer coupled with a hemispherical analyzer, under ultrahigh-vacuum conditions and with an Al Kα (hv=1486.74 eV) X-Ray source was used for the measurements acquisition. The measurements were performed at the ICN2 Photoemission Spectroscopy Facility. Charge effects were removed by taking the C1s line from adventitious carbon at 284.6 eV. For data analysis, CasaXPS software was used.

### X-ray Diffraction

200 µL of the sample was drop casted on a Si holder and dried in the oven at 50 °C. The spectra were collected in the 2θ scan range from 5° to 60° with a diffractometer (Malvern PANalytical X’Pert Pro MPD). For the measurements performed at the ICN2 XRD Facility, the x-ray source of a ceramic X-ray tube with Cu Kα anode (λ=1.540 Å) and the x’Celerator solid-state detector were used. The spectra were analyzed with X’Pert HighScore (version 2.2c (2.2.3)) software.

### Stability evaluation of GO:R848 complex

For the evaluation of long-term colloidal stability the purified GO:R848 complex and GO control, suspensions were stored at room temperature and in dark conditions. At specific time points, their pH was measured, their colloidal properties were assessed by DLS, and the R848 detachment from the GO surface was investigated.

### pH measurements

For the pH measurements, the FiveEasy™ FP20 pH meter equipped with the Mettler Toledo™ pH Electrode InLab Ultra-Micro-ISM was used.

### ζ-potential and hydrodynamic diameter measurements

The ζ-potential and hydrodynamic diameter were measured with a Zeta-sizer Nano ZS (Malvern Instruments) at the ICN2 Molecular Spectroscopy and Optical Microscopy Facility. 1 mL of samples at a GO concentration of 20 µg/mL were prepared and loaded in disposable capillary cells. The water dispersant settings for viscosity and refractive index were selected and each sample was measured three times at room temperature. The data were analyzed with Zetasizer (version 7.12) software and plotted as mean ± standard deviation unless stated otherwise.

### R848 detachment from GO surface

Drug detachment experiments for the purified complex were performed in specific time points by centrifuging 4 times at 4000 G, 20 °C for 10 minutes, using 100 kDa Amicon Ultra Centrifugal Filter devices. Similar to the purification process, UV-Vis spectroscopy and HPLC were used to determine the amount of R848 collected in each filtrate.

### Stability in cell culture medium

GO and GO:R848 were diluted in Dulbecco’s modified eagle medium (DMEM) supplemented with 1% Penicillin/Streptomycin and 10% fetal bovine serum (FBS) at a final concentration of 0.1 mg/mL and stored at room temperature. Visual colloidal properties were monitored at 0, 4, 8 and 24 h. Also, AFM analysis was performed at 0 and 8 h, where no colloidal agglomeration had started visually.

### Bone marrow-derived macrophage cell culture (BMDMs)

BMDMs were isolated from fibula and tibias of C57/BL6 female mouse and being filtered through a 100 µm cell strainer. Bone marrow cells were then cultured with Dulbecco’s modified eagle medium (DMEM) supplemented with 1% L-Glutamine, 1% Penicillin/Streptomycin, 10% fetal bovine serum (FBS) and 10 ng/mL murine-colony stimulating factor (M-CSF) (Peprotech, UK). Cells were cultured in non-treated cell culture dishes (Corning, UK), with 5% CO_2_, at 37 °C. Medium was refreshed on day 2, 4 and 6 with medium contained M-CSF at a final concentration of 10 ng/mL. Adherent cells were collected post-maturation period, on day 6-7 of differentiation for the *in vitro* experiments.

### Flow cytometry

BMDMs were plated in non-treated 24-well plates (Corning, UK) at a ratio of 200,000 cells/well. In parallel, cells were treated with LPS (100 ng/mL) / IFNgamma (20 ng/mL), GO (10 µg/mL), GO: R848 (10 µg/mL: 4 µg/mL) and R848 (4 µg/mL) and incubated for 24 hours with 5% CO_2_, at 37 °C. Cells were detached using 10mM EDTA for 10 min at 4°C and harvested for flow cytometry. BMDMs washed with PBS by centrifugation at 300 G, for 5 minutes at 8 °C and stained with Zombie UV, live/dead (BioLegend, USA) at 1:2000 dilution in PBS, for 15 minutes at room temperature. Cells were incubated with the conjugated primary antibodies F4/80-APC (1:200), CD11b-BV710 (1:100), CD80-FITC (1:100) and Fc receptor blocker (1:100) for 1 hour. Then cells were washed twice with flow buffer (PBS, 2mM EDTA, 2% FBS) at 500 G for 3 minutes at 8 °C and fixed with 1% PFA for 10 minutes at RT. Finally, cells were resuspended in 200 μL flow buffer and stored in the dark at 4 °C until analysis by flow cytometry. Flow cytometry was performed using MCCIR FCF BD LSR Fortessa (BD Bioscience, UK). Flow cytometry analysis was performed using FlowJo software (v10.6.1).

### ELISA

BMDMs were plated and treated as described above. Supernatants were collected and TNF-α ELISA MAX Deluxe Kit (BioLegend, UK), was used to performed ELISA according to manufacturer’s protocol.

### Live cell imaging

BMDMs were seeded at a density of 100,000/well in CellView 35mm, 4 compartment dishes (Griener BioOne, UK) and incubated with the treatments stated above for 2 hours. Post-treatment, cell supernatant was replaced with DMEM with CellMask green plasma membrane stain (Invitrogen, UK) and live cells were imaged under 5% CO_2_ at 37 °C, using confocal microscope 710 (ZEISS) with a Primo Plan-ACHTOMAT 40X/oil lens. Images were captured using ZEN 2010 B SP1 software using 594 and 405 lasers with gain of 1115 and 1012, respectively. Microscope settings were kept constant throughout the experiment. ZEN light software (version 9.1.2) was used for the image analysis. Bright field image and GO fluorescence signal images were compared to determine GO uptake and distinguish it from cell auto fluorescence.

### Statistical analysis

Data analysis and graphical design were performed using GraphPad Prism software (version 6.01). P-values were calculated using two-way ANOVA with Tukey’s post-hoc test for multiple comparisons. P-values of <0.05 were considered statistically significant. Data is plotted as mean ± standard deviation unless stated otherwise.

## Supporting Information

Supporting Information is available from the Wiley Online Library or from the author.

## Acknowledgments

The authors would like to thank the European Union Horizon 2020 Research and Innovation Programme under Grant Agreement no. 881603 (Graphene Flagship Core 3) to financially support this project. The ICN2 has been supported by the Severo Ochoa Centres of Excellence programme [SEV-2017-0706] and is currently supported by Grant CEX2021-001214-S funded by MCIN/AEI/10.13039.501100011033. The authors would also like to acknowledge the United Kingdom Research and Innovation (UKRI) Engineering and Physical Sciences Research Council (EPSRC) 2D-Health Programme Grant (EP/P00119X/1). The authors are also thanking Angeliki Karakasidi for the synthesis and characterization of the GO material. Lastly, the authors are acknowledging the ICN2 Advanced Electronic Materials and Devices Group, headed by Prof. Jose A. Garrido for the AFM and Raman measurements that were performed in their laboratory.

## Author contributions

D.D performed the main studies including protocol optimization for the complex preparation, physicochemical characterizations and colloidal stability studies. M.A. assisted with the physicochemical characterizations. M.S. and T.K. conducted the in vitro experiments and data interpretation. K.K. conceptualized this study and D.D., N.L. and T.K. contributed in the experimental design. K.K. and N.L. coordinated the experimental work. All authors contributed to the manuscript’s writing.

## Supporting Information

**Figure S1.**
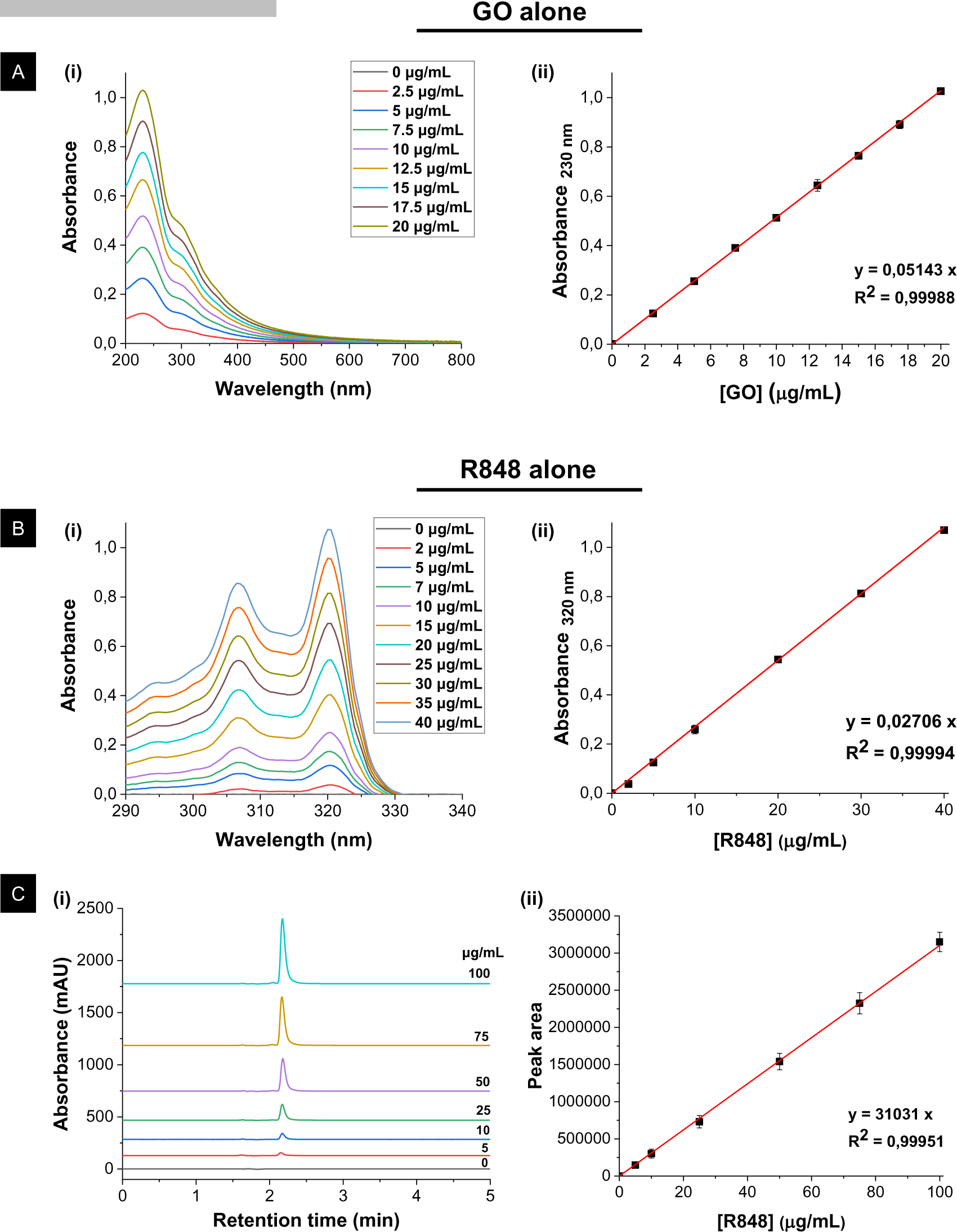
Calibration data of GO and R848 alone. **(A)** UV-Vis calibration data of GO at concentrations ranging between 0 and 20 μg/mL at the wavelength of 230 nm. **(B)** UV-Vis calibration curve of R848 at concentrations ranging between 0 and 40 μg/mL at the wavelength of 320 nm. **(C)** HPLC calibration curve of R848 at concentrations ranging between 0 and 100 μg/mL at the wavelength of 254 nm. In each case, the following is presented: **(i)** raw data of one indicative experimental run; **(ii)** calibration curves from data expressed as mean±SD (n=3).

**Figure S2.**
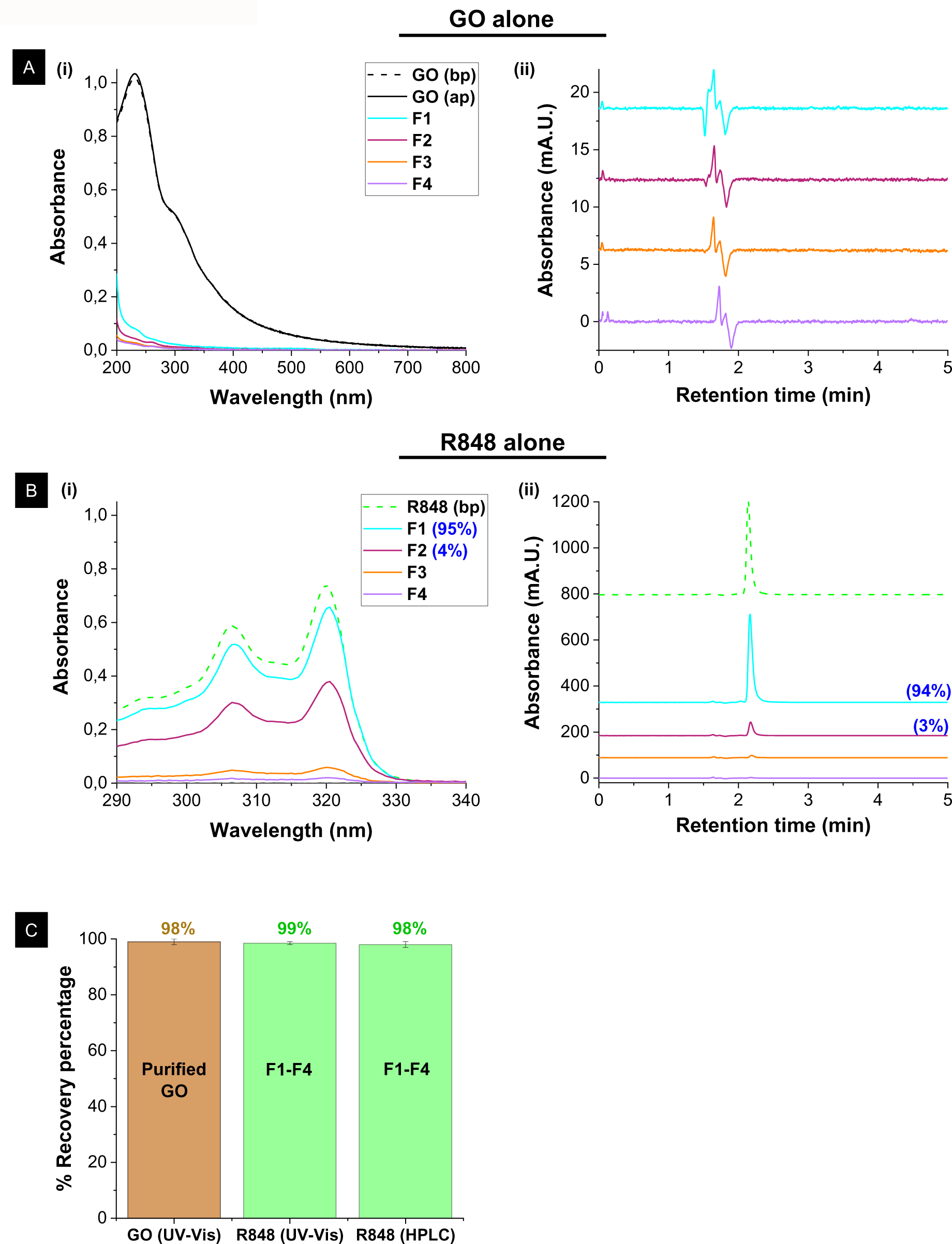
Validation of purification method for GO and R848 controls. **(A)** GO control purification. **(i)** UV-Vis spectrum before (bp) (dashed line) and after (ap) purification (solid line) (GO (20 µg/mL)), and signal of the corresponding filtrates (F1-F4) in the range 200-800 nm; **(ii)** HPLC chromatogram of GO filtrates (F1-F4). **(B)** R848 control purification. **(i)** UV-Vis spectrum before purification (bp) (dashed green line) and signal of the corresponding filtrates (F1-F4) in the range of 290-340 nm**; (ii)** HPLC chromatogram of R848 before purification (bp) and of the corresponding filtrates (F1-F4). All R848 quantity at the end of the procedure is present only in the filtrates. **(C)** Percentage (%) GO recovery in the purified sample fraction on the filter (brown bar); and Percentage (%) R848 recovery washed in the filtrates (green bars) quantified by the two independent techniques and measurements: UV-Vis and HPLC spectroscopy. The results (generated by applying the formula (1)) are expressed as a mean±SD of n=3 replicates.

**Figure S3.**
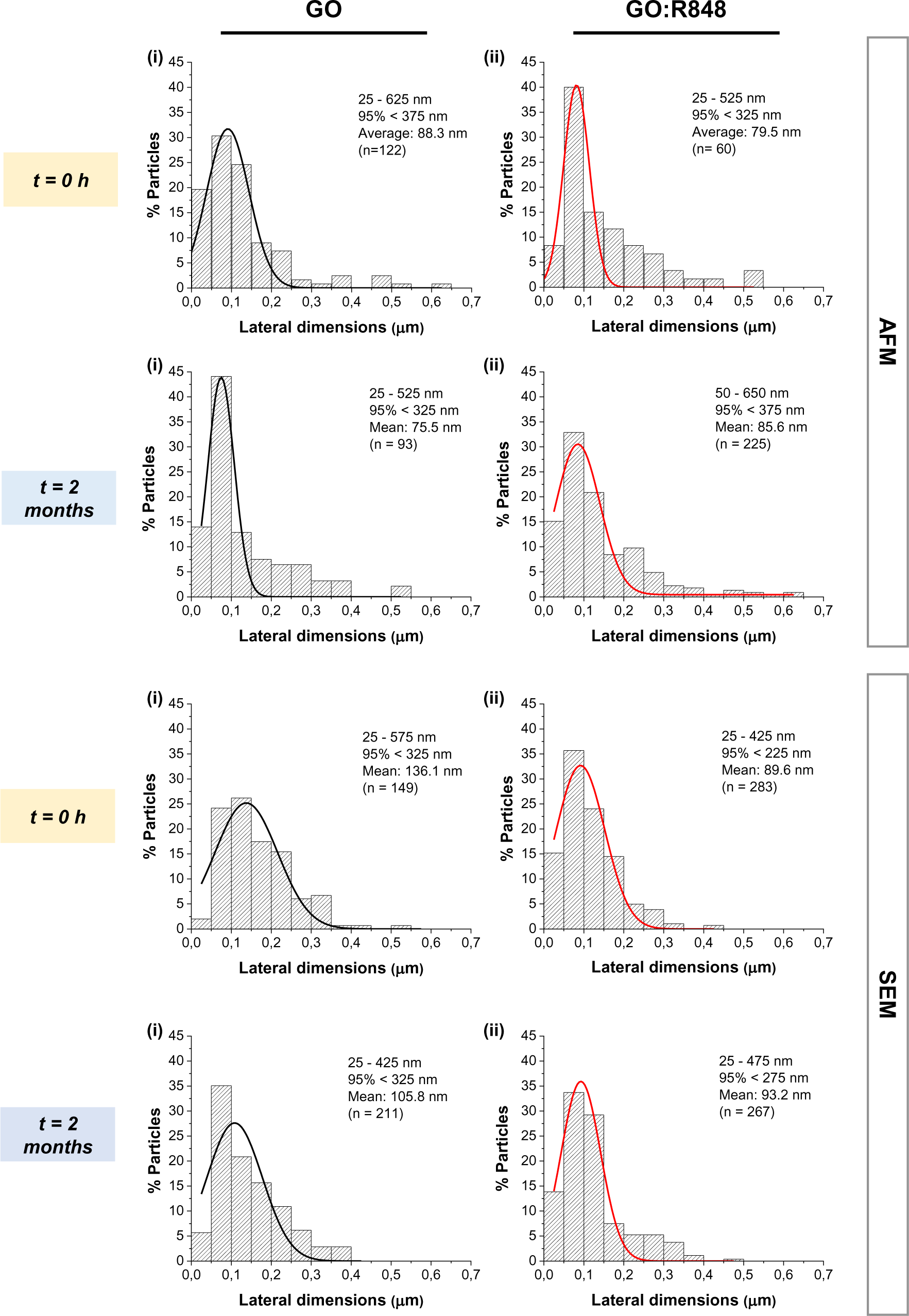
Size distribution of GO nanosheets alone and in complex by AFM and SEM. The data was obtained from AFM and SEM images at t=0 h (yellow) and t=2 months (blue) for **(i)** GO control; **(ii)** GO:R848 complex after purification (ap) complex.

**Figure S4.**
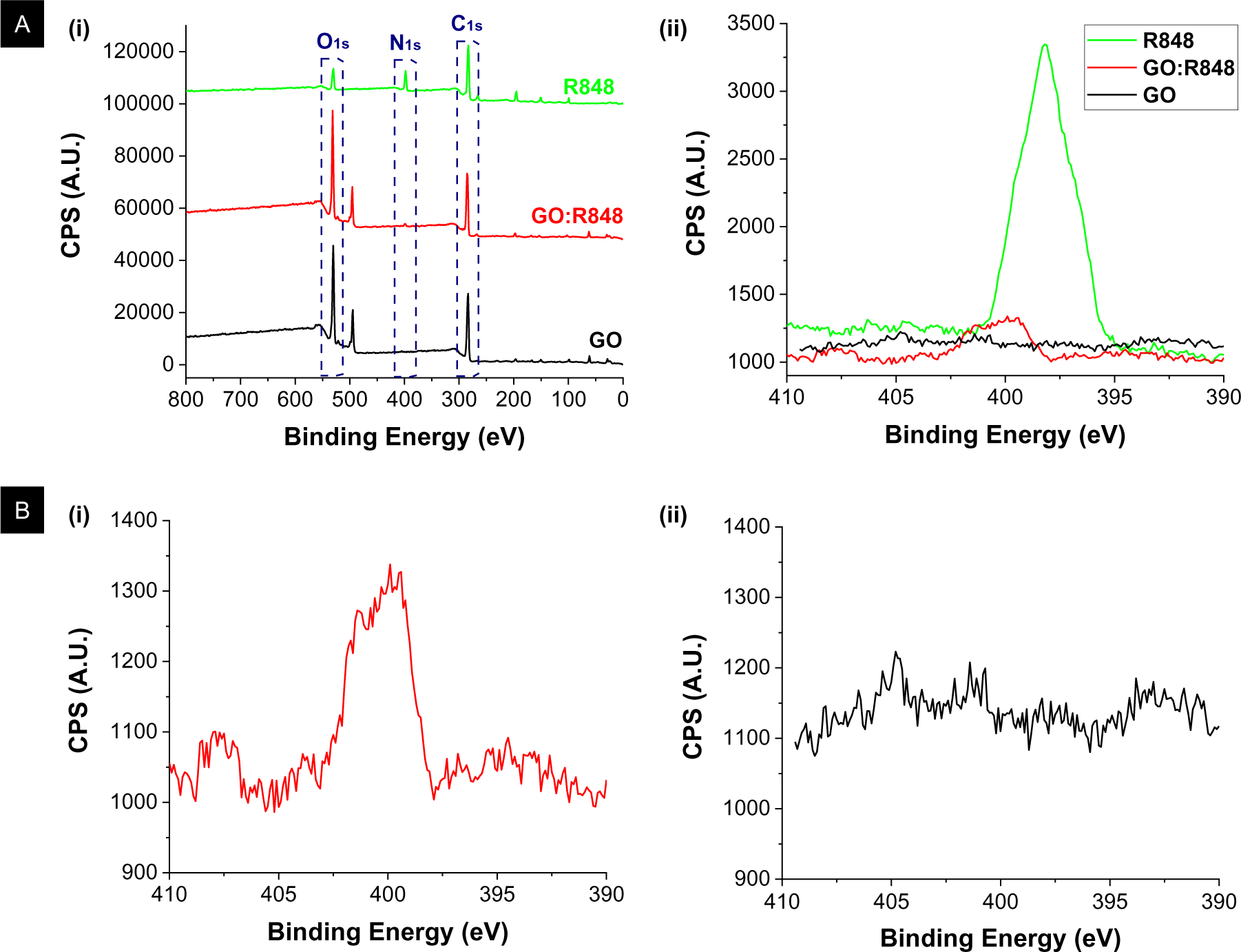
XPS analysis of GO:R848 complex after purification (ap). **(A)** XPS high-resolution spectra; **(i)** Characteristic peaks of C, O, and N highlighted for GO:R848 complex (red), GO (black) and R848 (green) controls in the range 800-0 eV; **(ii)** N_1s_ high-resolution spectra in the range 410-390 eV (CPS is counts per seconds). **(B)** Magnification of N_1s_ high-resolution spectra. **(i)** GO:R848 complex; **(ii)** GO control.

**Figure S5.**
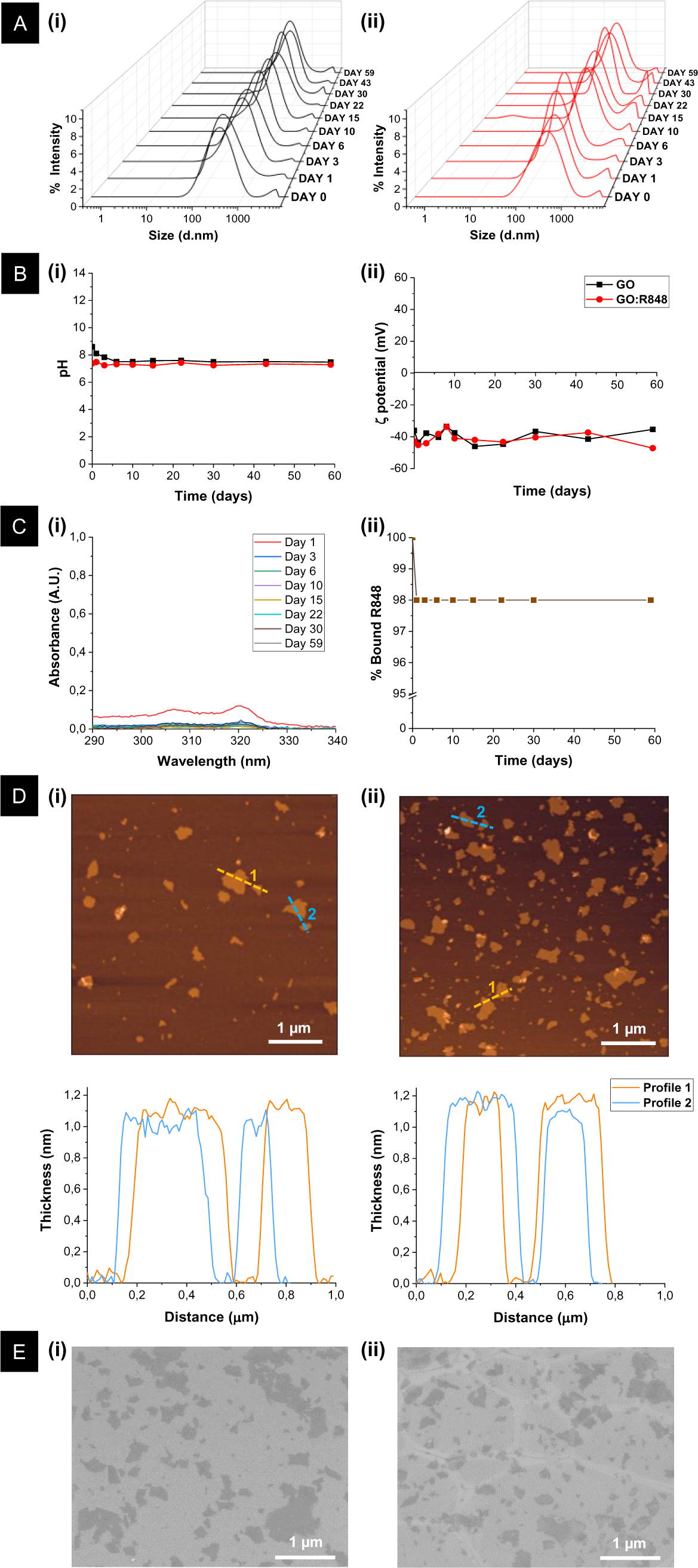
Colloidal stability of GO:R848 complex after purification (ap) over 2 months. **(A)** Dynamic light scattering (DLS) plots showing the evolution of mean particle size over time. **(i)** GO control; **(ii)** GO:R848 complex. **(B)** Measurements over 2 months. **(i)** pH monitoring; **(ii)** Average particle surface charge (ζ-potential) for GO control (black) and GO:R848 complex (red). **(C) (i)** UV-Vis spectrum of GO:R848 filtrates (unbound R848) in the range of 290-340 nm over 2 months; Note that the spectrum shown in each time point is the sum of absorbance from all individual filtrates (F1-F4); **(ii)** Percentage (%) of R848 that remains bound on the GO surface over 2 months. **(D)** AFM height images after 2 months from corresponding nanosheets cross section and thickness graphs (shown underneath) for **(i)** GO control; **(ii)** GO:R848 complex. **(E)** SEM micrographs after 2 months for **(i)** GO control; **(ii)** GO:R848 complex. In all images scale bars are 1μm.

**Figure S6.**
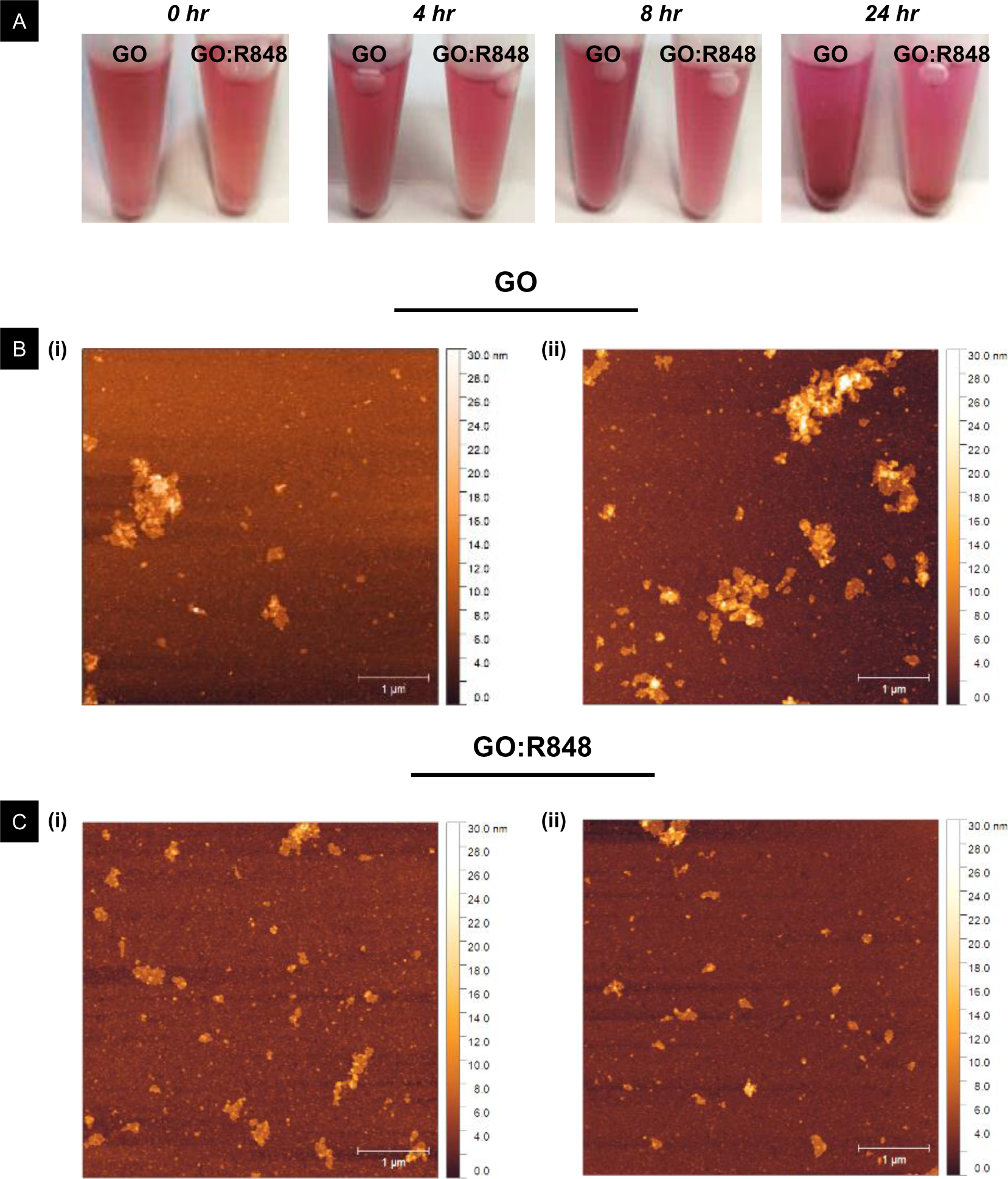
Colloidal stability of GO:R848 complexes after purification (ap) in cell culture media (DMEM + 10% FBS) for 24 hours. **(A)** Visual aspect of GO control and GO:R848 complex in media at various time points: 0, 4, 8 and 24 hours. **(B)** AFM height images of GO control for **(i)** 0 hours and **(ii)** 8 hours. **(C)** AFM height images of GO:R848 complex for **(i)** 0 hours and **(ii)** 8 hours. Note that all samples were measured at a final GO concentration of 0.1mg/mL after dilution in media.

**Figure S7.**
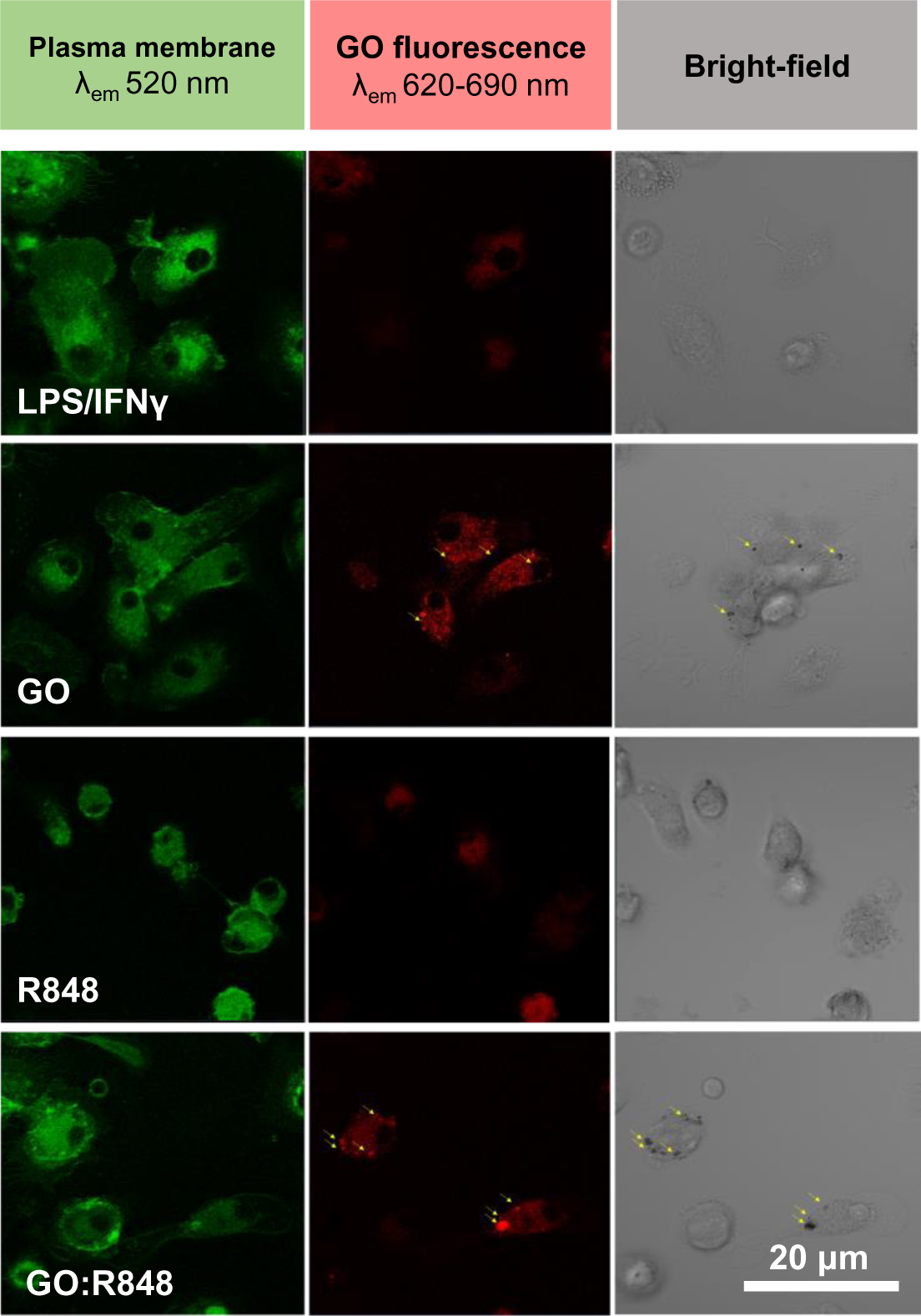
Confocal images taken with zoom factor 1, showing the co-localisation of GO and GO:R848 (10:4) with BMDMs membrane dye (green), 2hr post treatment. Red fluorescence field showing the auto-fluorescence of GO. Bright field confirmed the presence of GO black particles. Images were taken via confocal microscope at 40x magnification. Representative images from n=6/condition, Scale bar 20 μm.

**Figure S8.**
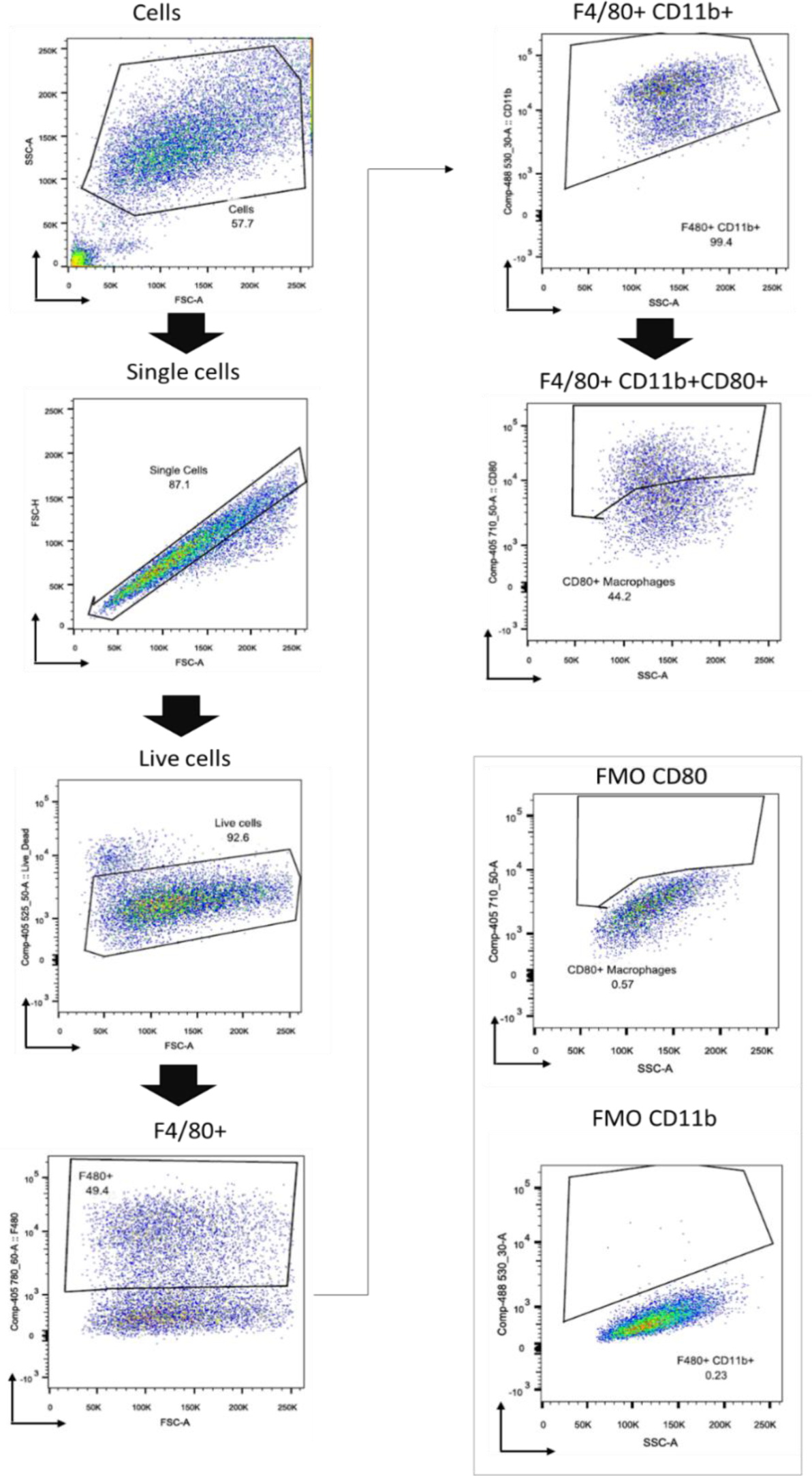
Flow cytometry dot plots and gating strategy used for CD80+ / F480+ CD11b+ cells. Indicative FACS data is shown of cells first isolated from debris; single cells gated to avoid clumps; live cells selected to avoid dead cells. F480+ cells were taken forward from live cells and based on the fluorescence minus one (FMO) controls of CD11b and CD80, CD11b+ and CD80+ cells were gated.

**Table S1.** Main physicochemical properties of GO nanosheets, used in the present study. Properties including lateral dimensions, thickness, optical properties, degree of defects, interlayer distance, surface charge, functionalization degree, chemical composition, and purity, are indicated below.

| Material property | Technique | Result |
| --- | --- | --- |
| Lateral dimensions | SEM | 50 - 950 nm [n = 946] |
|  |  | 95% < 450 nm |
|  |  | Mean 144 nm |
|  | AFM | 10 - 890 nm [n = 2890] |
|  |  | 95% < 350 nm |
|  |  | Mean 72 nm |
| Thickness | AFM | 1 nm (1 layer) |
| Optical properties | Absorption spectroscopy | $\epsilon_{232}$ (mL $\mu\text{g}^{-1}$ cm$^{-1}$)= 0.05 |
| Degree of defects ($I_D/I_G$) | Raman spectroscopy | 1.12 ± 0.02 [n=3] |
| Peak (2θ) | XRD | 12.36 ° [n=1] |
| Interlayer distance (nm) |  | 0.71 |
| Surface charge (ζ-Potential) | Electrophoretic mobility | - 47.2 ± 0.7 mV |
| Functional groups | FTIR | $\nu(\text{C-H})$: 2838 cm$^{-1}$ |
| | | $\nu(\text{C=O})$: 1735 cm$^{-1}$ |
| | | $\nu(\text{C=C})$: 1645 cm$^{-1}$ |
| | | $\nu(\text{O-H})$: 1423 cm$^{-1}$ |
| | | $\nu(\text{C-O})$: 1062 cm$^{-1}$ |
| Functionalization degree | TGA | 30-75°C: 7% (water) |
|  |  | 200-250°C: 30% |
|  |  | 250-950°C: 20% |
|  |  | TOTAL 50% |
| Chemical composition (%) | XPS | O: 29.6, C: 68.5, N: 1.1, S: 0.9 |
| Purity % (C+O) |  | 98,0 |
| C:O ratio |  | 2,3 |
| π-$\pi^*$, O-C=O, C=O, C-O, C=C (%) | | 1.4, 7.0, 3.1, 44.3, 44.3 |

